# Punctuated evolution of myxoma virus: rapid and disjunct evolution of a recent viral lineage in Australia

**DOI:** 10.1101/465583

**Authors:** Peter J. Kerr, John-Sebastian Eden, Francesca Di Giallonardo, David Peacock, June Liu, Tanja Strive, Andrew F. Read, Edward C. Holmes

## Abstract

Myxoma virus (MYXV) has been evolving in a novel host species – European rabbits – in Australia since 1950. Previous studies of viruses sampled from 1950 to 1999 revealed a remarkably clock-like evolutionary process across all Australian lineages of MYXV. Through an analysis of 49 newly generated MYXV genome sequences isolated in Australia between 2008 and 2017 we show that MYXV evolution in Australia can be characterized by three lineages, one of which exhibited a greatly elevated rate of evolutionary change and a dramatic break-down of temporal structure. Phylogenetic analysis revealed that this apparently punctuated evolutionary event occurred between 1996 and 2012. The branch leading to the rapidly evolving lineage contained a relatively high number of non-synonymous substitutions, and viruses in this lineage reversed a mutation found in the progenitor standard laboratory strain (SLS) and all previous sequences that disrupts the reading frame of the *M005L/R* gene. Analysis of genes encoding proteins involved in DNA synthesis or RNA transcription did not reveal any mutations likely to cause rapid evolution. Although there was some evidence for recombination across the MYXV phylogeny, this was not associated with the increase in evolutionary rate. The period from 1996 to 2012 saw significant declines in wild rabbit numbers, due to the introduction of rabbit hemorrhagic disease and prolonged drought in south-eastern Australia, followed by the partial recovery of populations. We therefore suggest that a rapidly changing environment for virus transmission changed the selection pressures faced by MYXV and altered the course of virus evolution.

**IMPORTANCE:** The co-evolution of myxoma virus (MYXV) and European rabbits in Australia is one of the most important natural ‘experiments’ in evolutionary biology, providing insights into virus adaptation to new hosts and the evolution of virulence. Previous studies of MYXV evolution have also shown that the virus evolves both relatively rapidly and in a strongly clock-like manner. Using newly acquired MYXV genome sequences from Australia we show that the virus has experienced a dramatic change in evolutionary behavior over the last 20 years, with a break-down in clock-like structure, the appearance of a rapidly evolving virus lineage, and the accumulation of multiple non-synonymous and indel mutations. We suggest that this punctuated evolutionary event likely reflects a change in selection pressures as rabbit numbers declined following the introduction of rabbit hemorrhagic disease virus and drought in the geographic regions inhabited by rabbits.

---

An experimental field study in Australia in 1950 to test whether *Myxoma virus* (MYXV) could be used as a biological control for European rabbits (*Oryctolagus cuniculus*) accidentally initiated one of the largest and best known natural experiments in host-pathogen coevolution. Detailed studies on the coevolution of the virus with its new host over the subsequent 60 years have done much to inform thinking on pathogen evolution in general and the evolution of virulence in particular (1-5).

MYXV is a poxvirus of the *Leporipoxvirus* genus (family *Poxviridae*; subfamily *Chordopoxvirinae*). The natural host of the virus released in Australia is the South American tapeti (forest rabbit: *Sylvilagus minensis*) (6), also called *Sylvilagus brasiliensis* (7, 8). In the tapeti, MYXV causes a localized, cutaneous fibroma at the inoculation site (6). Virus is passively transmitted on the mouthparts of biting insects such as mosquitoes or fleas probing through the virus-rich fibroma. In general, MYXV does not appear to cause significant clinical disease in the natural host. However, when European rabbits (*Oryctolagus cuniculus*) are infected the virus spreads systemically, overwhelming the immune system, to cause the lethal, disseminated disease myxomatosis (9). Related viruses, *Rabbit fibroma virus* and Californian MYXV, occur in separate *Sylvilagus* spp in North America.

Following its experimental release, MYXV unexpectedly spread across large swathes of rabbit-infested south-eastern Australia (10). The released virus, originally isolated in Brazil and later termed the Standard Laboratory Strain (SLS), had a case fatality rate (CFR) estimated at 99.8% (11). However, there was rapid natural selection for slightly attenuated viruses which, by allowing longer survival of the infected rabbit, were more efficiently transmitted by the mosquito vectors (12-13).

To study the evolution of virulence field isolates of MYXV were classified into five virulence grades based on CFR and average survival time (AST) in small groups of laboratory rabbits, which are the same species and had essentially the same outcomes following infection as wild European rabbits. Moderately attenuated grade 3 strains, with CFRs of 70-95%, predominated in the field in the decades following the release. More attenuated grade 4 viruses were less common, while highly virulent grade 1 and 2 viruses became relatively rare despite ongoing releases of virulent virus. Highly attenuated grade 5 viruses with CFRs <50% were never common (2, 13-16). At the same time there was a strong selection for resistance to myxomatosis in the wild rabbit population. The short generation interval of rabbits meant that this evolutionary process could be measured in real-time by challenging successive generations of rabbits with virus of known virulence. At one study site, CFRs dropped from 90% to 26% over a 7 year period (17-18).

The deliberate release in France of a separate Brazilian isolate of MYXV in 1952 led to the establishment and spread of MYXV in Europe. Later termed the Lausanne strain (Lu), this virus was more virulent than SLS (2, 19). However, although the starting virus, ecological conditions and vectors were different, the evolutionary outcomes in Europe at the phenotypic level were remarkably similar to those in Australia, with the emergence of moderately attenuated strains of virus and selection for genetic resistance in the wild rabbit population (16). The Lu sequence is the type sequence for MYXV. This virus was originally isolated in 1949, and possesses a genome of 161,777 bp of linear double stranded DNA with inverted terminal repeats (TIRs) of 11,577 bp and closed single strand hairpin loops at the termini. The genome encodes 158 unique open reading frames (ORFs), 12 of which are duplicated in the TIRs (20, 21).

The virus originally released in Australia, SLS, is believed to be derived from an isolate made in Brazil around 1910 (22). A sample of this virus was later transferred to the Rockefeller Institute (23) where it was maintained by passage in laboratory rabbits. MYXV used for testing in Australia and Britain was obtained from the Rockefeller in the 1930s (24). SLS has a genome of 161,763 bp with the same organization as Lu, but with single base indels disrupting two ORFs *M005L/R* in the TIRs and *M083L* (4). Although the Lu strain of MYXV was widely released in Australia for tactical rabbit control from the 1970s to 1990s, there is no evidence that this virus has established (25, 26) and all of the viruses we have sequenced are derived from SLS.

Detailed molecular studies, in particular gene knock-out experiments using the Lu strain, have revealed much about the way in which MYXV subverts and overwhelms the immune system of the European rabbit (9, 27). In many cases, deletion of a single “virulence gene” is sufficient to substantially attenuate the virus. Sequencing studies of field isolates of MYXV from both the Australian and European radiations have shown that reading frame disruptions in key virulence genes are not uncommon (4, 28-30). However, characterization of some of these viruses has revealed that disruption to virulence genes can be compatible with high virulence, and that recent viruses from both radiations exhibit a novel immune suppressive phenotype in laboratory rabbits (5, 30). In addition, reverse genetics to correct potentially attenuating mutations or insert a potentially attenuating mutation did not result in phenotypic changes (31). This has made it difficult to define the molecular basis of the initial attenuation or to define the molecular drivers of the ongoing virus evolution in wild rabbit populations, potentially in competition with continuing selection for resistance to myxomatosis.

Phylogenetic analyses of almost 50 full genome sequences from Australian and European viruses have demonstrated that despite the similarity in evolutionary outcomes on the two continents there were almost no shared mutations across the Australian and UK sequences and hence no convergent evolution at the genomic level (30). However, the evolutionary rates for the two radiations were both remarkably similar, at approximately 1 × 10^−5^ nucleotide substitutions per site, per year. Virus evolution on both continents was also strongly clock-like, manifest in a consistent relationship between time of sampling and genetic distance (30).

Previous genomic analyses for the Australian viruses only included viruses sampled from 1950 to 1999, with most samples obtained between 1990 and 1996. However, the ecological environment facing rabbits in Australia changed dramatically in the mid-1990s with the release of a second biocontrol agent – *Rabbit hemorrhagic disease virus* (RHDV) – that greatly reduced the size of the rabbit population (32-35). To determine whether MYXV evolution differs markedly before and after the emergence of RHDV, we sequenced a large set MYXV isolates sampled between 2008 and 2016, extending previous studies by nearly 20 years and considerably expanding geographic coverage. In addition, to investigate possible variation in the progenitor virus, we sequenced SLS stocks prepared in 1949 and 1956.

## RESULTS

### Phylogenetic analysis of Australian MYXV

We sequenced and analyzed 50 field strains of MYXV sampled in south-eastern Australia. Of these, 49 were collected during the period 2008-2016. One archival sample from 1990 was also sequenced (Table 1). In addition, we re-sequenced the SLS progenitor and an SLS-like virus from archival samples, and re-sequenced regions of the Glenfield (GV) strain (originally sampled in February 1951 during the first epizootic). The geographic locations of the sampled viruses, as well as their lineage (see below) are shown in Fig. 1.

**Table 1.**
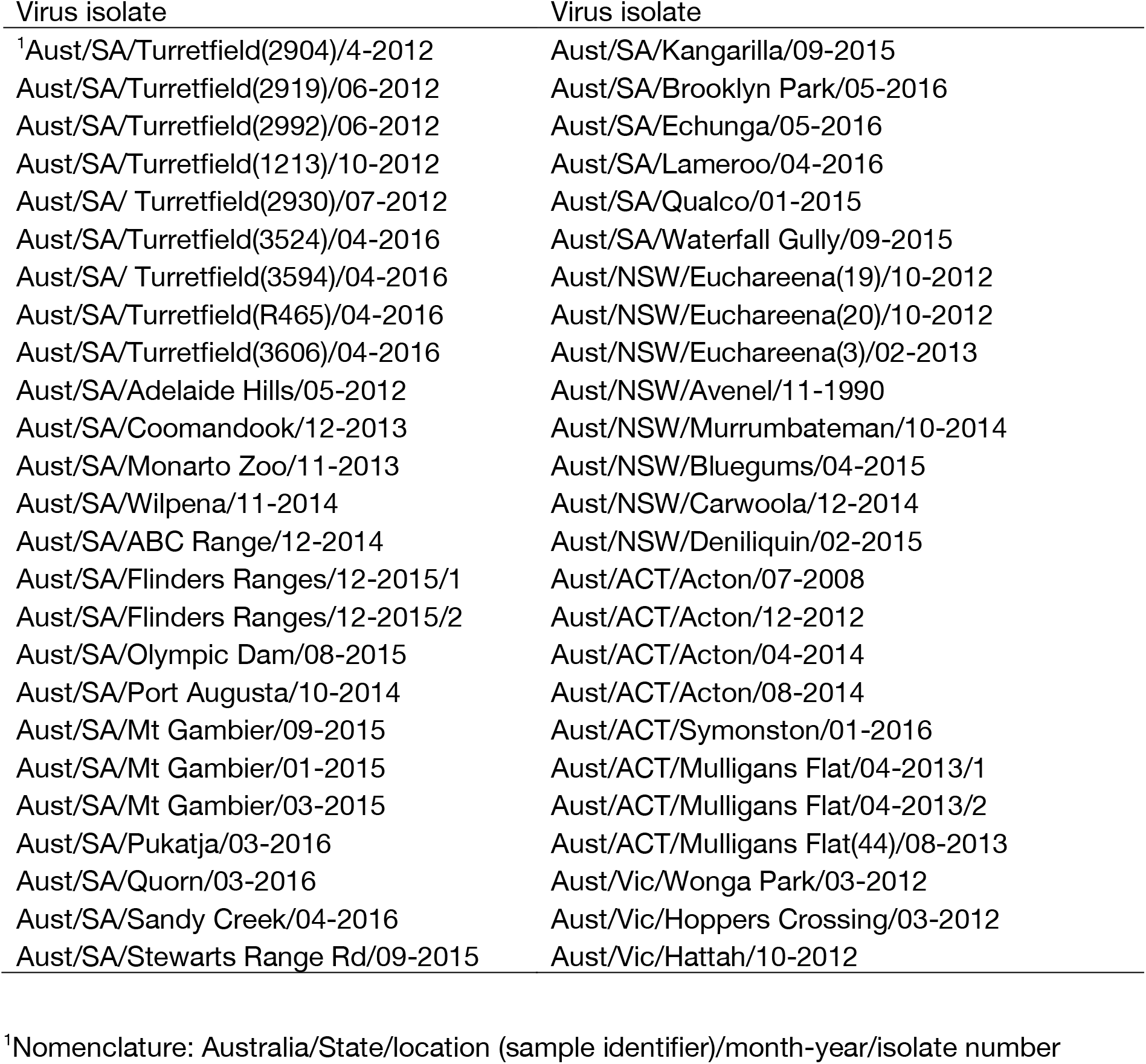
Australian field isolates of MYXV sequenced as part of this study.

**Fig. 1.**
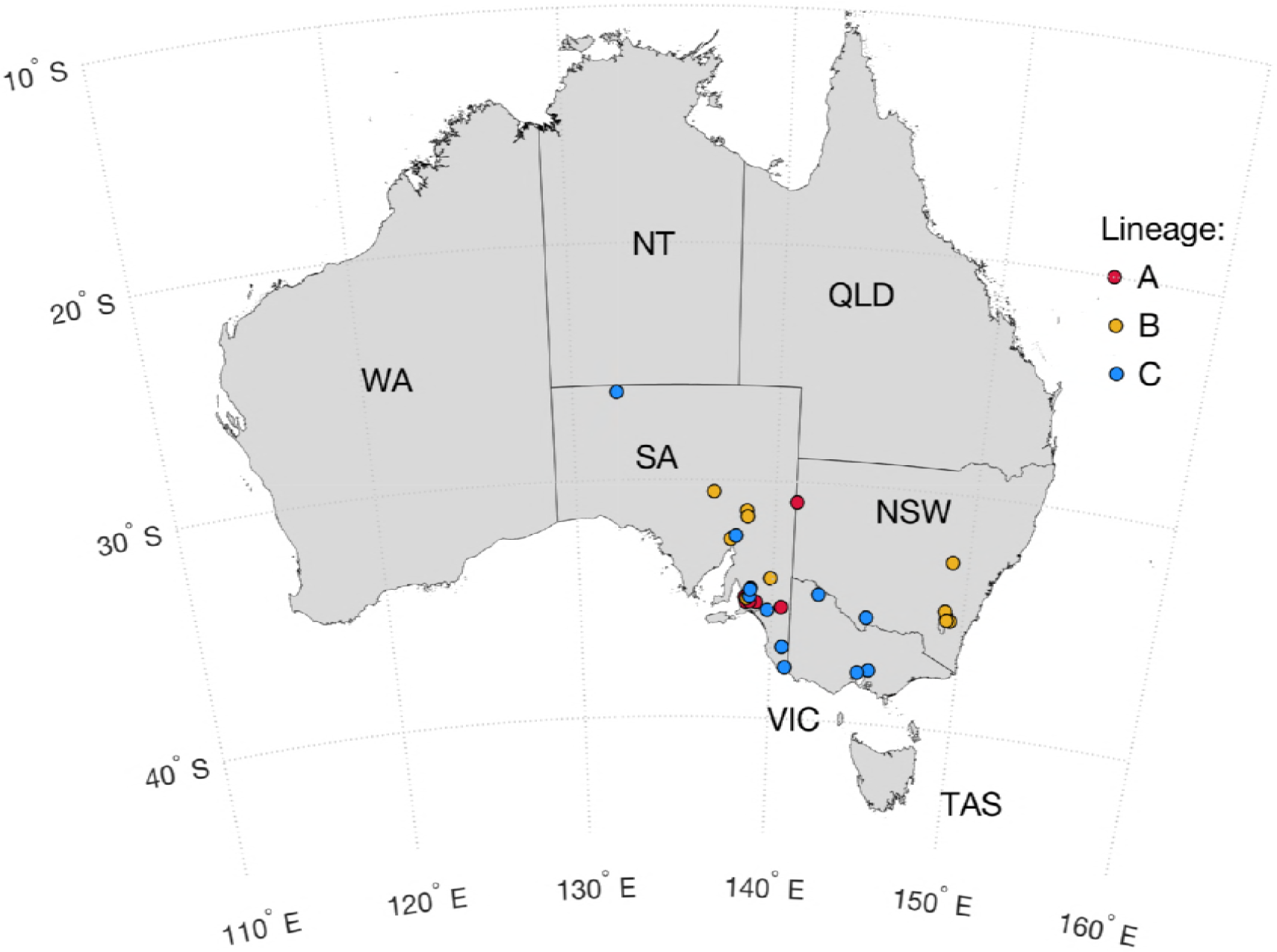
Sampling locations in Australia of the new sequenced isolates of MYXV. The sampling locations are color-coded by phylogenetic lineage (see Results and Fig. 2). The different Australian states and Territories are marked.

Our phylogenetic analysis of all complete MYXV genomes from Australia (n = 78) revealed that those viruses sampled between 1990-2016 can be placed into three main lineages, denoted here as lineages A, B and C, although only lineage C receives strong bootstrap support (Fig. 2). A single virus, Meby, isolated in Tasmania in 1991 could not be placed in these three lineages and fell in a more basal position. Lineage A contained viruses sampled between 1990-2016, including a small cluster of five viruses isolated between 2013 and 2016 (Fig. 2). Similarly, lineage B contained viruses sampled from 1993-2016 and from a variety of geographic and climatic areas including semi-arid South Australia (Pt. Augusta, Olympic Dam, Flinders Ranges, Wilpena, ABC Range) and more temperate, higher rainfall areas in South Australia, New South Wales and close to Canberra in the Australian Capital Territory (Fig. 1 and 2). This lineage also included previously sequenced viruses isolated from the Canberra region and western Queensland in the 1990s. Interestingly, all but four viruses in clade B, including all the recent isolates, have a 1635 nt duplication encompassing the *M156R*, *M154L* genes and part of the *M153R* genes normally found at the RH TIR boundary that has been duplicated at the LH TIR boundary; this is effectively an expansion of the TIRs, with loss of 923 nt of the 3’ end of the *M009L* gene at the LH TIR (Fig. 2). This duplication was first seen in viruses isolated from the Canberra district in 1995 (26, 28).

**Fig. 2.**
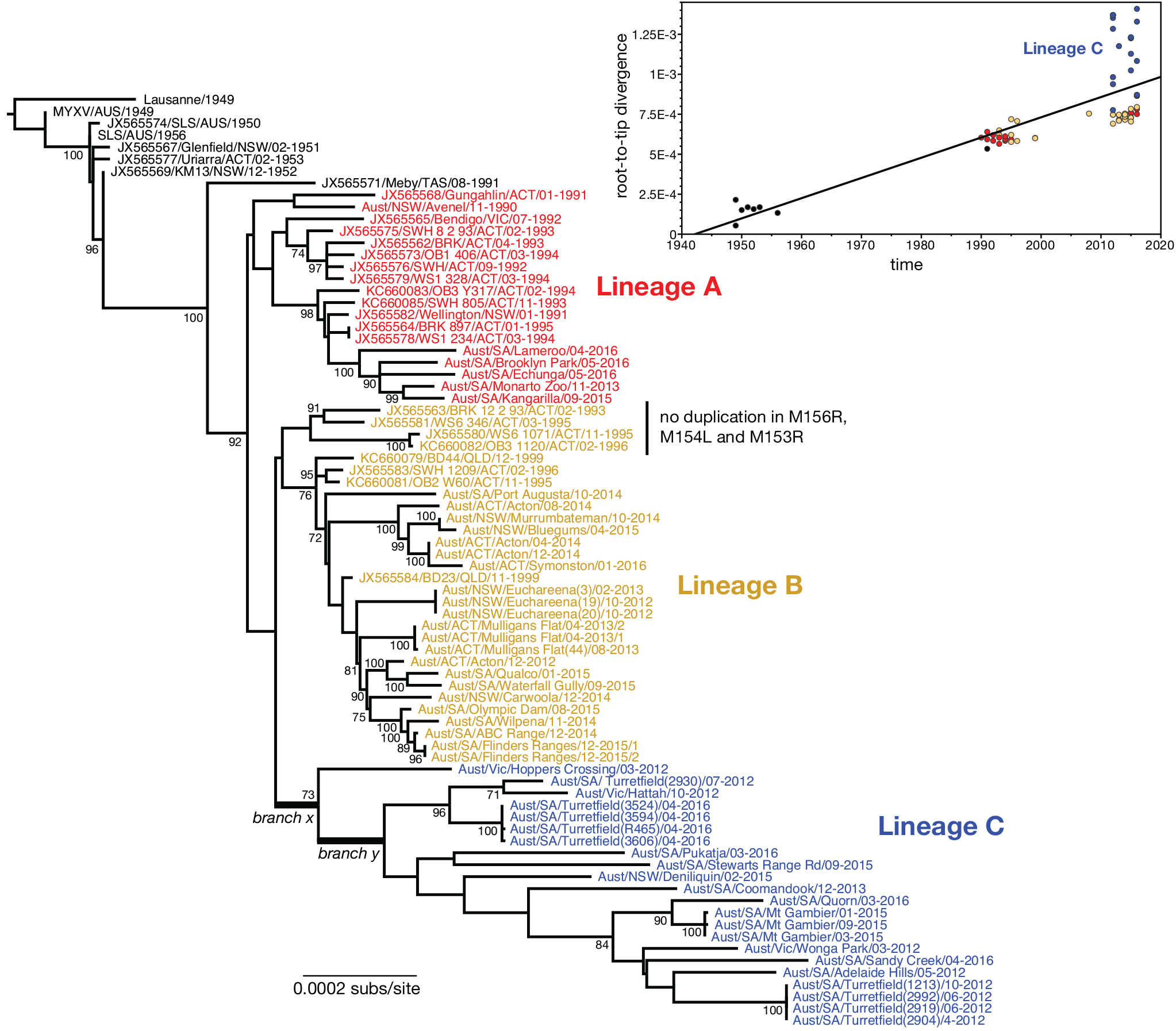
Evolutionary history of MYXV in Australia. The tree depicts the phylogenetic relationships among all 78 complete genome sequences of MYXV sequenced to date. All viruses sampled from 1990 have been color-coded, and with exception of the Meby strain sample in Tasmania in 1991 fall into three main lineages (denoted A, B and C). Two branches – denoted x and y – associated with the emergence of the rapidly evolving lineage C are indicated. All viruses within lineages B and C aside from the four marked possess a duplication in *M156R*, *M154L* and part of *M153R*. The tree is rooted by the European Lausanne strain, and all branches are scaled according to the number of nucleotide substitutions per site. All bootstrap values >70% are shown. Insert: regression of root-to-tip genetic distances against year of sampling for all available Australian isolates of MYXV. Each point is color-coded according to their phylogenetic lineage. The recently sampled lineage C, which has experienced an elevation in evolutionary rate, is marked.

Finally, lineage C includes viruses isolated from different geographic and climatic areas of Victoria (Hattah, Hoppers Crossing, Wonga Park), southern NSW (Deniliquin) and a broad geographic and climatic region of South Australia from the far north desert country (Pukatja) to the higher rainfall Adelaide Hills and Mt. Gambier in the south. This lineage is notable in that it only contains viruses sampled from 2012 onwards and that all viruses contain the TIR expansion seen in clade B. More dramatically, individual tree branches within lineage C are longer than those in the other two lineages and exhibit a marked lack of temporal structure; that is, many of the earlier sampled viruses (from 2012) fall no closer to the root of the tree than those sampled in 2016. This includes viruses sampled between 2012 and 2016 at the same site – Turretfield in South Australia. That viruses sampled in 2012 occupy phylogenetic positions both at the root and toward the tips of this lineage suggests that most of the divergence in this lineage, and hence the rapid evolutionary event, had occurred prior to 2012.

To assess the extent of temporal structure in the Australian MYXV sequences in a more quantitative manner we performed a regression of root-to-tip genetic distances against the year of sampling. This revealed that all MYXV sequences with the exception of those assigned to lineage C exhibited relatively strong rate constancy, as previously observed in MYXV in both Australia and Europe (30). In contrast, most of those viruses assigned to lineage C have elevated rates such that they consistently fall above the regression line (Figure 2 insert). Across the Australian viruses as a whole, the R^2^ value of the relationship between root-to-tip genetic distances against the year of sampling was 0.58, whereas values of between 0.93 and 0.98 were observed in previous studies that did not include lineage C. Importantly, the viruses assigned to lineage C were sequenced at the same time as those from the other lineages indicating that their divergent position is not a laboratory artefact.

### Mutations associated with the rapidly evolving MYXV lineage C

To determine what processes might be associated with the rate elevation in lineage C, we mapped all nucleotide and amino acid substitutions onto the ML phylogenetic tree and examined in detail those that occurred early in the history of lineage C. Specifically, we focused on (i) the branch directly leading to lineage C, immediately prior to the divergence of the Hopper’s Crossing virus (designed branch x; Fig. 2), and (ii) the first (i.e. next) internal branch in lineage C (designated branch y).

A total 10 nucleotide substitutions were reconstructed as falling on branch x, nine of which were non-synonymous. Such a relatively high frequency of non-synonymous substitution is compatible with either a relaxation of selective constraints or, more likely, the occurrence of positive selection on this lineage. These non-synonymous substitutions include an amino acid change in the *M010L* gene (A66V). M010 is a homologue of cellular epidermal growth factor/transforming growth factor alpha and is critical for virulence (36, 37). Interestingly, this is the only amino acid substitution in M010 we have seen in Australian isolates, although a valine is also found at this position in the Californian MYXV strain MSW and rabbit fibroma virus. Two non-synonymous mutations were also seen in *M036L* that encodes a protein orthologous to the orthopoxvirus *Vaccinia virus* (VACV) O1. O1 potentiates ERK1/2 signalling stimulated by the VACV epidermal growth factor homologue (38). There are no data on the role (if any) of M036 in MYXV infection and whether it potentiates the activity of M010. However, mutations in the *M036L* gene are relatively common and disruption of the gene in field isolates of MYXV is compatible with high virulence (5, 25, 30).

In addition to the novel mutations found on this branch there were two reversals of non-synonymous mutations in *M014L* and *M134R* that occurred on the root branch leading to all the sequences from 1990 onwards. The role of M014 is undefined, but it likely functions as part of an E3 Ub ligase complex (39). M134 is an orthologue of *Variola virus* (VARV) B22, which has been implicated in suppression of T cells in orthopoxvirus infections (40).

A total of 19 substitutions were constructed to fall on branch y (Figure 2), 10 of which are non-synonymous and six represent reversal of mutations occurring on branches leading to all the post-1990 sequences. However, perhaps of most interest is an additional single nucleotide deletion of a G nucleotide at position 6353 at nt 35 in the 5’ end of the *M005L* ORF (and in the corresponding position in *M005R*) that corrects a G insert at this position found in SLS and in all subsequent Australian virus sequences previously sequenced. The G insert disrupts the *M005L/R* ORF with stop codons after codon 78 and an altered amino acid sequence after codon 11. Targeted disruption of the *M005L/R* gene in Lu caused severe attenuation of the virus with case fatality rates reduced from 100% to 0 (41). All of the viruses within rapidly evolving lineage C have an intact *M005L/R* ORF with the exception of Hoppers Crossing.

### Recombination in Australian MYXV

Recombination may be relatively common in poxviruses (42) and has been observed between MYXV and rabbit fibroma virus (43, 44) and between South American and Californian MYXV (45) (46). Despite this, we previously found no evidence for recombination in either the Australian or European MYXV (4, 28, 30).

An analysis of possible recombination events in our expanded data of Australian MYXV revealed a complex pattern of phylogenetic movement that is compatible with the action of recombination, although the location of precise recombination breakpoints is difficult to determine. An analysis of recombination events using a suite of methods within the RDP package failed to identify either consistent (i.e. statistically significant) recombination events or recombination breakpoints. For example, although some analyses suggest that Hattah (2012), Coomandook (2013) and Adelaide Hills (2012) are recombinant, this is not confirmed in more detailed phylogenetic analyses because there is insufficient bootstrap support to confirm phylogenetic incongruence (data not shown).

In an additional attempt to reveal recombinants we inferred phylogenetic trees for non-overlapping 25K regions of the MYXV genome (Fig. 3). This revealed clear, although complex, patterns of topological movement, and in most cases it was impossible to statistically prove recombination because of a lack of phylogenetic resolution due to low levels of sequence diversity. Indeed, the only case in which the changes in tree topology were supported by high levels of bootstrap support concerned a cluster of three viruses from Euchareena sampled in NSW in 2012 and 2013 and a singleton comprising a virus from Port Augusta in South Australia sampled in 2014. These viruses fell into lineage B in phylogenies of the whole virus genome, as well as that of region 25-50K, but clustered with some lineage C viruses in the region 75-100K, particularly four viruses sampled at Turretfield (South Australia) in 2016 (Fig. 3).

**Fig. 3.**
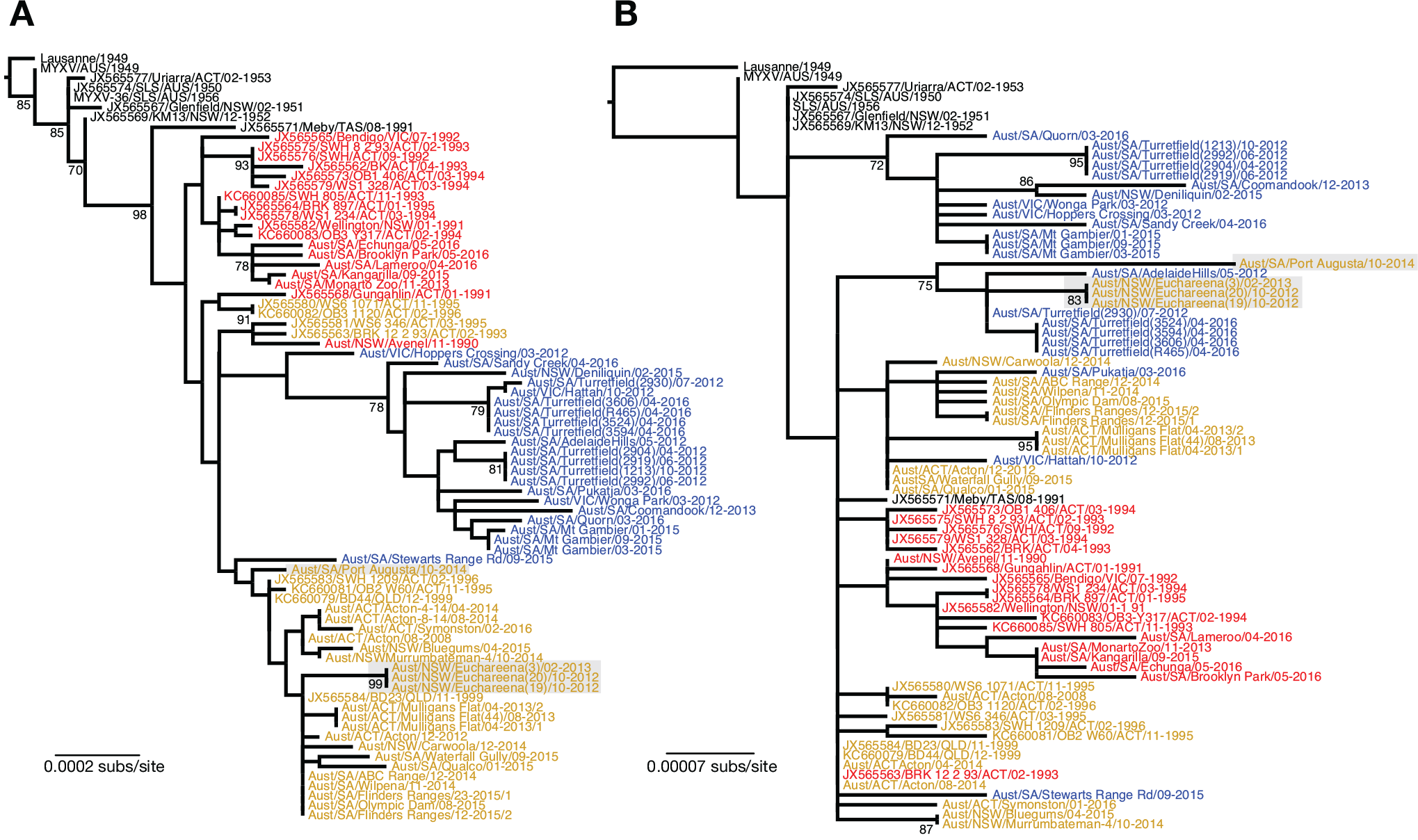
Phylogenetic evidence for recombination in Australian MYXV. Tree (A) is for the aligned genomic region 25-50K while tree (B) comprises the region 75-100K. All taxa are color-coded according to their whole genome lineage (see Fig. 2). Putative recombinant strains are shaded. All nodes with bootstrap support values >70% are shown. All trees are drawn to a scale of nucleotide substitution per site and rooted by the Lausanne sequence.

In addition, as noted below, some individual mutations exhibit phylogenetic distributions that are compatible with the occurrence of recombination, although this cannot be demonstrated in a statistically robust manner.

### Gene-specific mutations in Australian MYXV analysis

Across all Australian MYXV sequenced to date, 13 genes have no mutations and 16 have only synonymous mutations. These conserved genes largely encode structural proteins, fusion/entry proteins or those involved in genome replication or transcription. The exceptions include *M128L* that encodes an immunomodulatory protein important for virulence (47), and *M123R*, an orthologue of VACV *A35R*, that encodes an immunomodulatory protein (48, 49). Multiple genes have in-frame indels, reading frame disrupting indels or stop codons (Table 2).

**Table 2.**
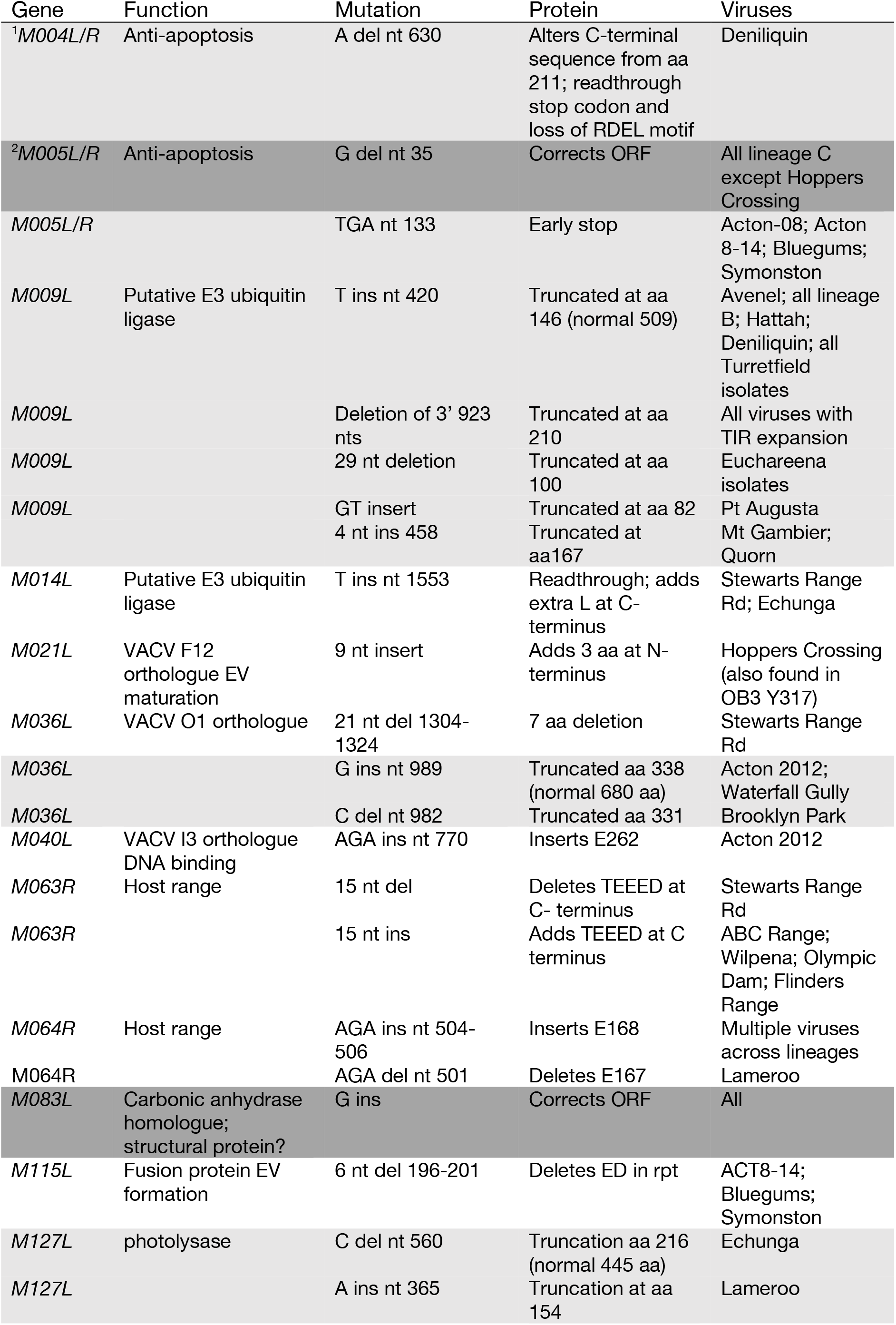
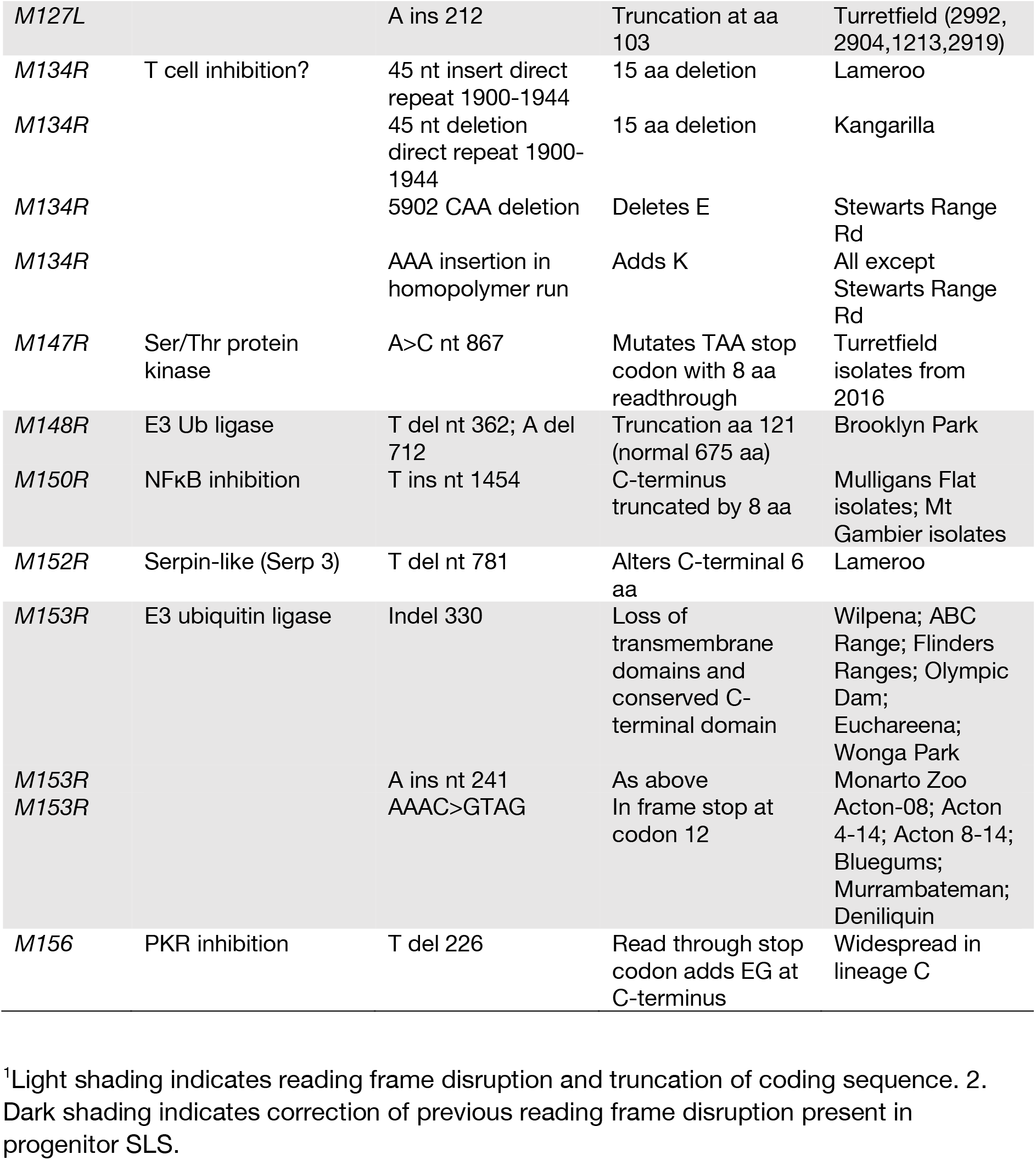
Insertions, deletions, disruptions and repairs in genes from new MYXV sequences.

#### Mutations in proteins involved in DNA replication and transcription

We examined whether high evolutionary rates could have been due to a mutator phenotype or possibly more rapid replication. The MYXV DNA polymerase complex consists of four proteins: M034 (DNA pol; VACV E9), M079 (DNA processivity/uracil DNA glycosylase; VACV D4), M080 (helicase/primase; VACV D5), and M111 (DNA pol processivity; VACV A20) (50). M034 and M079 had no amino acid substitutions in any virus. A single amino acid substitution was observed in M080 in one virus (Bluegums), and another amino acid change, R430C in M111, was identified in some of the rapidly evolving viruses from lineage C (Wonga Park, Adelaide Hills, Turretfield 2012 isolates, Mt. Gambier, Pukatja, Quorn, Sandy Creek). Published mutational studies of VACV A20 do not provide any information on the phenotypic impact of this substitution (51-53), although the cysteine at this position is also found in rabbit fibroma virus.

In addition to the polymerase complex, M112 (Holliday junction resolvase; VACV A22), M133 (DNA ligase; VACV A50) and M074 (DNA topoisomerase; VACV H6R) are required for DNA replication. None of the genes for these proteins contained any non-synonymous substitutions. Proteins involved in generating nucleotide precursors, M061 (thymidine kinase), M015 (ribonucleotide reductase small subunit), M012 (dUTPase), had no consistent mutations in lineage C viruses.

Of 27 genes encoding proteins involved in virus transcription, most had only synonymous mutations, mutations that occur only in single viruses, or mutations common to most of the Australian viruses. However, there were two mutations in M044, the viral NPH II protein (nucleoside triphosphatase/RNA helicase), that occur uniquely in most lineage C viruses (R118C; P478S). Of note was that a number of viruses (Wonga Park, Hattah, Adelaide Hills, Turretfield 2012 and 2016 isolates, Coomandook, Deniliquin, Mt Gambier, Pukatja and Sandy Creek) have P478S, although the proline at this site (within the helicase C-terminal domain) is otherwise completely conserved across the chordopoxviruses.

Interestingly, there is a widespread mutation in the C-terminus of the RNA polymerase subunit encoded by *M114R* (VACV A24) in Australian MYXV (P1147H). Many of the viruses with this M114 mutation also share a mutation in M156 (L71P), the MYXV member of the poxvirus K3 family of protein kinase R (PKR) inhibitors, that abolishes PKR inhibition (54). A T1120M mutation in the VACV RNA polymerase subunit A24 was able to confer resistance to PKR in an artificial evolution system (49).

#### Analysis of MYXV promoters

We examined mutations up to 100 nt 5’ to ATG start codons to determine whether there are lineage-specific differences in promoter sequences. Since only a small number of MYXV genes have had their promoters mapped, promoters have necessarily been inferred by homology with conserved chordopoxvirus early, intermediate and late promoter sequences (55). The majority (but not all) viruses in lineage C have a T deletion at the 3’ end of the *M156R* gene (Table 2), that forms part of the upstream T rich region of the *M008.1L/R* late promoter. This mutation is also seen in three lineage B viruses, and might be expected to decrease transcription from this promoter (56). An additional A at the 5’ end of the *M040L* E promoter sequence is present in most recent Australian isolates in lineages A and B, but not lineage C, which might be expected to have a minor increase in transcription efficacy (57). Most other mutations were considered unlikely to have an effect on promoters or occurred only in single sequences.

### Mutations causing loss of the *M153R* ORF are common and persist in the field

The *M153R* gene encodes a 206 amino acid RING-CH E3 ubiquitin ligase that downregulates MHC-1, CD4, Fas/CD95 and ALCAM/CD166 from the surface of infected cells *in vitro* (58-61). Targeted deletion of this gene in Lu causes marked attenuation of the virus (58). However, mutations disrupting the ORF and predicted to lead to loss of function are common in Australian isolates of MXYV.

Our previous sampling of field isolates of Australian MYXV did not allow us to determine whether any of the viruses with disrupted *M153R* genes left descendants. However, with increased sampling, we can now identify multiple viruses with *M153R* disruptions and their descendants. The first of these was initially seen in a lineage B virus isolated in Canberra in 2008 (Acton 07-2008), and in later isolates sampled from the same site (Acton 04-2014, Acton 08-2014), ∼35 km north (Murrumbateman 2014 and Bluegums 2015), and 10 km to the south-east (Symonston 2016). This is a complex mutation in which three nucleotide substitutions within a four nucleotide span, AAAC>GTAG, form an in-frame TAG stop codon after codon 11. The next in-frame ATG is at nt 275. The identical mutation is found in a lineage C virus isolated at Deniliquin in 2015, some 400 km west of Canberra. The complex nature of this mutation makes it unlikely to have occurred more than once and its presence in viruses in different lineages may therefore reflect the occurrence of recombination.

In the second example, a G insertion at nt 330, causing disruption of the *M153R* ORF with an early stop after amino acid 124, was found in viruses isolated from Wilpena and ABC Range in the Flinders Ranges of outback South Australia in November and December 2014. The same mutation was present in related viruses isolated 12 months later in the Flinders Ranges and in August 2015 at Olympic Dam some 100 km to the west. While we cannot exclude the possibility that this insertion occurred multiple times independently, it seems more likely that these closely related viruses were descendants of a single common ancestral strain that contained the mutation.

### Sequences of the Australian progenitor SLS virus

The original epizootic of myxomatosis in 1950 stemmed from wild rabbits inoculated with virus later termed Standard Laboratory Strain (SLS). In the following years, many releases of this virus were made by inoculating rabbits. It is possible that polymorphisms in stocks contributed to diversity in the field. For example, the nucleotides at positions 2576 and 149,127 in the stock of SLS that we originally sequenced were not found in any other Australian viruses. This suggests that mutations at these positions may have occurred during passage in rabbits or cell culture (4) or existed as polymorphisms in stocks of SLS used for release. To test this, we grew and sequenced virus from stocks originally obtained from Professor Frank Fenner and labelled as “SLS (1956)” and “Myxoma virus 1949”.

There are a number of sequence differences between our original SLS sequence and the 1956 stock and 1949 viruses (Table 3). In the case of nt positions 2756 and 149,127 the SLS 1956 stock had the same sequence as the field isolates, suggesting that these were the original sequences at these positions. In addition, the 1956 virus had two single nt insertions which were seen in some early isolates: the G insertion at 122,397 was present in the GV strain (isolated in February 1951, but more virulent than SLS despite the gene disruptions), and the A insertion at 131,587 was present in the Ur strain (1953). Hence, the originally released SLS may have comprised a mixture of sequences. The provenance of the 1949 virus is not known. It is clearly closely related to SLS and distinct from Lu. Fenner obtained and tested virus from the Rockefeller Institute (13) from which SLS was originally derived. If this is that virus, then it appears that SLS has diverged during passage. In addition, using a new stock of virus, we confirmed gene disruptions in *M014L*, *M130R* and *M153R* originally identified in the high virulence GV strain (4).

**Table 3.**
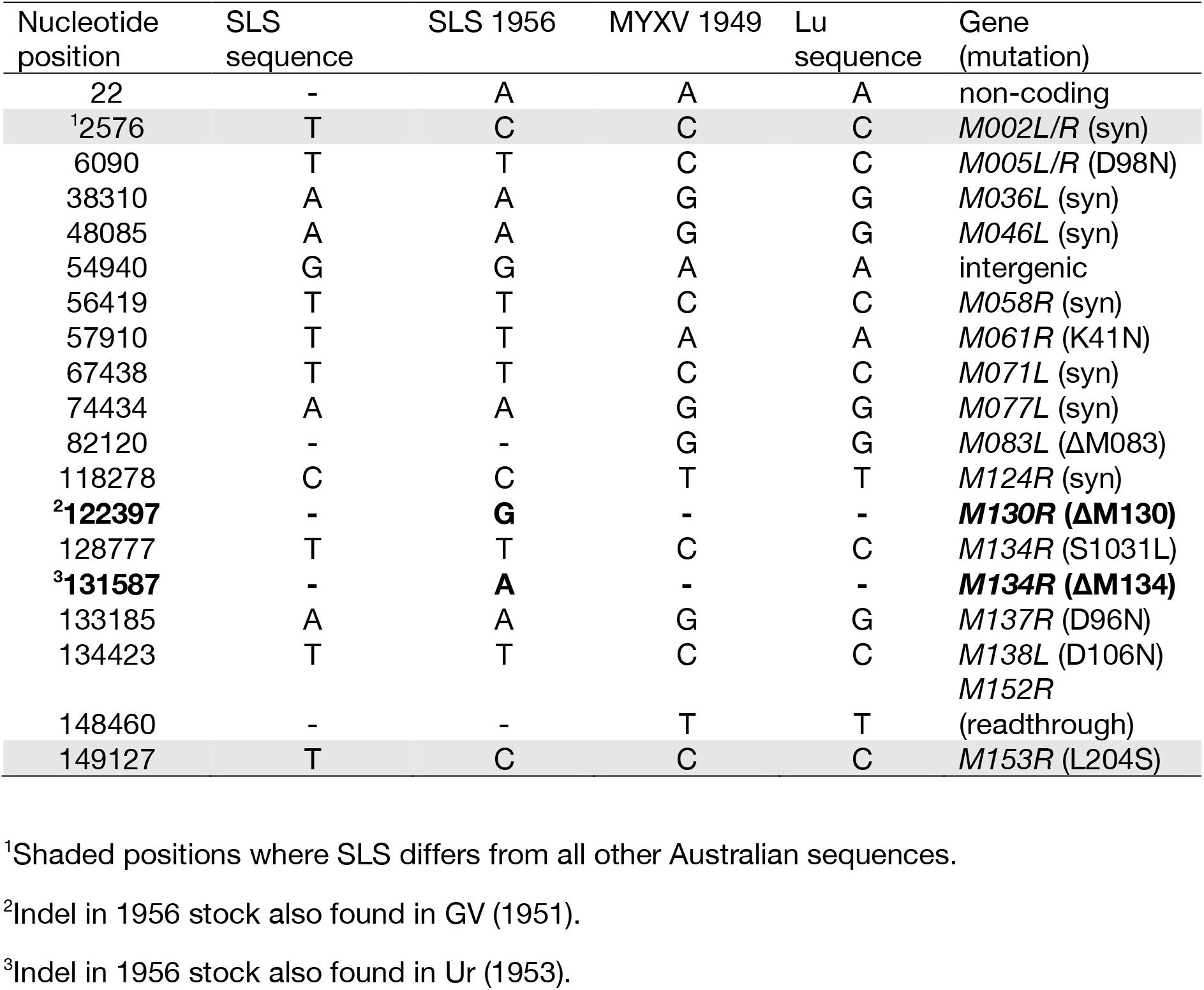
Sequence variation in SLS stocks.

| Nucleotide position | SLS sequence | SLS 1956 | MYXV 1949 | Lu sequence | Gene (mutation) |
| --- | --- | --- | --- | --- | --- |
| 22 | - | A | A | A | non-coding |
| <sup>1</sup>2576 | T | C | C | C | *M002L/R* (syn) |
| 6090 | T | T | C | C | *M005L/R* (D98N) |
| 38310 | A | A | G | G | *M036L* (syn) |
| 48085 | A | A | G | G | *M046L* (syn) |
| 54940 | G | G | A | A | intergenic |
| 56419 | T | T | C | C | *M058R* (syn) |
| 57910 | T | T | A | A | *M061R* (K41N) |
| 67438 | T | T | C | C | *M071L* (syn) |
| 74434 | A | A | G | G | *M077L* (syn) |
| 82120 | - | - | G | G | *M083L* (ΔM083) |
| 118278 | C | C | T | T | *M124R* (syn) |
| <sup>2</sup>122397 | - | **G** | - | - | ***M130R* (ΔM130)** |
| 128777 | T | T | C | C | *M134R* (S1031L) |
| <sup>3</sup>131587 | - | **A** | - | - | ***M134R* (ΔM134)** |
| 133185 | A | A | G | G | *M137R* (D96N) |
| 134423 | T | T | C | C | *M138L* (D106N) |
| 148460 | - | - | T | T | *M152R* (readthrough) |
| 149127 | T | C | C | C | *M153R* (L204S) |

<sup>1</sup>Shaded positions where SLS differs from all other Australian sequences.
<sup>2</sup>Indel in 1956 stock also found in GV (1951).
<sup>3</sup>Indel in 1956 stock also found in Ur (1953).

## DISCUSSION

Previous studies of MYXV evolution had shown that the evolutionary rate in this virus since its release in Australia and Europe was comparatively high for a DNA virus, at ∼1 × 10^−5^ nucleotide substitutions per site, per year (4, 30) – and extremely clock-like in both continents (30). While this still appears to be the case for more recent isolates assigned to lineages A and B, it is clear that a third and large group of viruses, designated as lineage C and that fall in diverse locations across south-eastern Australia, have much greater genetic diversity and a break-down of clock-like structure, compatible with a greatly elevated rate of evolutionary change. The temporal structure in lineage C, with the earliest sampled viruses from 2012 falling at diverse phylogenetic locations including both close to the root and near the tips, suggests that the full diversity in this lineage, and hence the rate elevation, was established at some point between 1996 (i.e. the most recent date of the majority of our last sample of sequences) and 2012.

What, then, is the most likely cause of this punctuated evolutionary event? We did not find any convincing evidence for mutations in genes involved in DNA replication or transcription that might lead to a more error-prone replication, or more cycles of replication, that could have conceivably led to an increase in rate. Similarly, while there is some evidence for recombination in the recent isolates of MYXV, including some viruses within lineage C, its impact appears to be localized to specific viruses and particular genomic locations. We therefore suspect that recombination is unlikely to be responsible for the punctuated evolution of lineage C.

If changes in mutation and replication dynamics, as well as the occurrence of recombination, can be discounted, then a change in selection pressures during the period 1996-2012 seems the most likely explanation for the emergence of the divergent viruses associated with lineage C. Two major changes in rabbit populations occurred during this period and these could conceivably have influenced transmission of MYXV and hence altered selection pressures. From 1995, a novel pathogen, RHDV, spread through Australia causing rabbit population crashes in many areas (32-35, 63, 64), reducing the number of MYXV-susceptible hosts. In addition, a major, prolonged drought occurred over south-eastern Australia between 1996 and 2009 putting additional pressure on rabbit populations. A possible third influence is the establishment of the Spanish rabbit flea (*Xenopyslla cunicularis*) as a vector for MYXV in the arid areas of south-eastern Australia, although there is little information on its impact. Decreased mosquito numbers due to dry conditions may have meant that fleas became the major vectors for transmission of MYXV, in turn making transmission much more localized or completely failing in regions where fleas were absent. Changes in the timing of epizootics of myxomatosis were also seen (63). Smaller, fragmented rabbit populations may have favored viruses with longer average persistence in the individual rabbit and local extinctions of MYXV were likely commonplace. As rabbit populations recovered from both the initial impact of rabbit hemorrhagic disease (66) and drought, any viruses better able persist, perhaps by adaptation to more efficient transmission, would have had a selective advantage.

Within lineage C it is tempting to speculate that the reversion of the *M005L/R* gene mutation to generate an intact ORF provided the phenotype conferring that selective advantage. The M005 protein has been shown to inhibit apoptosis in MYXV-infected lymphocytes in European rabbits and to have host-range functions in human cancer cells, and is a key virulence gene in Lu (41, 67, 68). Mutational disruption of M005 has been shown to reduce virulence in the Lu genetic background with no development of secondary cutaneous lesions (41). Australian isolates do not appear to have expressed M005 despite many having high virulence. The role of virulence in transmission in low density rabbit populations with high genetic resistance is likely complex and it is possible that mutations acting on other phenotypic characteristics such as lesion morphology may be important. The genetic determinants of virulence in the field are clearly complex and likely multifactorial, and wild-type virulence is not easily explained by the so-called virulence determinants identified in gene knock-out studies.

Aside from the potential selection pressures acting on lineage C, there is the surprising success of viruses with the duplication of *M154L* and *M156R*. Whether there is a selective advantage in duplication of two potential immunomodulatory genes or loss of the *M009L* gene is not clear. M154 has homology to VACV M2, which inhibits NF-κB activity (69), while M156 inhibits PKR (70). In artificial evolution experiments using VACV, duplication of PKR inhibitors such as K3L occurred quickly and provided a selective advantage (71, 72, 71). Interestingly, a mutation (L71P) in *M156R* that is present in all of the viruses in lineage B and in the duplicated *M156R* genes has been demonstrated to inactivate PKR inhibition (54). Despite this loss of function, some viruses with this mutation have high virulence in laboratory rabbits (5, 25). In addition, reverse engineering a virulent virus to have the 71P mutation did not alter virulence (31). However, most viruses in lineage C, all of which have the duplication, do not have the L71P mutation indicating a reversion at this position. The majority of these viruses also have a T deletion in the 3’ end of the *M156R* gene causing readthrough of the stop codon and the addition of two amino acids (E76 and G77) at the C-terminus.

The *M009L* gene encodes a 509 amino acid protein that is likely part of an E3 ubiquitin ligase complex (39). *M009L* has suffered multiple disruptions during the evolution of MYXV in Australia (Table 2) (4). The *M009L* ORF was also lost in the Californian MSW strain of MYXV and in the Kazza strain of rabbit fibroma virus (46, 55). Only one group of viruses sequenced here has an intact *M009L* ORF - Kangarilla, Brooklyn Park, Lameroo, Echunga and Monarto in lineage A - as does the previously sequenced Bendigo virus from 1992 (also lineage A).

In sum, during the period 1996-2012, viruses with the *M154L*/*M156R* gene duplication and comprising the newly described lineages B and C have become dominant in our sampling over a wide geographic area of south-eastern Australia. Within this group is a highly divergent and rapidly evolving virus lineage in which the normally constant rate of molecular evolution has been disrupted. Although we are uncertain what evolutionary forces are responsible for this punctuated evolutionary event, we suggest that it most likely reflects a combination of the combined impact of RHDV and a large-scale drought, both of which would have severely reduced rabbit numbers and in turn changed the selective environment facing MYXV. Similarly, while it is currently unclear what mutations may have been most impacted by these selective events, the mutations in *M005L/R* and *M156R*, which are of plausible phenotypic importance, should clearly be a starting point in further investigations.

## MATERIALS AND METHODS

### Field samples for virus isolation

Samples for virus isolation were submitted either as tissues (usually eyelids) or conjunctival swabs from rabbits with clinical signs of myxomatosis found dead or collected for other purposes. All samples were processed and viruses isolated on RK13 cell monolayers as previously described (30).

### Archival samples of SLS and Glenfield virus (GV)

Small glass straws containing freeze dried aliquots of SLS produced in rabbits were originally obtained from the late Professor Frank Fenner (Australian National University) in 1988. A vial labelled "myxoma virus 1949" and containing freeze dried material was also present from the same source. Virus from these vials was amplified in RK13 cells. Vials of freeze dried Glenfield virus were produced by the Commonwealth Serum laboratories in 1962 as homogenates of tissue from infected rabbits.

### DNA preparation

Virus was concentrated from infected RK13 cells by osmotically bursting the cells, centrifugation to remove nuclei and other cell debris, DNase and RNase digestion followed by polyethylene glycol precipitation (30, 73, 74). Viral DNA was prepared using the Masterpure DNA extraction kit (Epicorp).

### PCR analysis and sequencing of disrupted genes in GV

The primer sequences used were as follows:

M153R: F: AATGGAGGTGTTATAAACGCGACTGCCACG 3’; R:
CGATCTTTGTTATAGACAAACGAGATACCT
M014L: F: GCCAGGTCTATTCTGTCGATC; R:
CGTCATTTACGTCTTGGGAGGAGTCTCGTAC
M130R: F: TATATTATACACGGCCTTATTCGACGGCG; R:
CGTCGTACGGAAGGTGACTGTCTACGTTAA

Sequencing of the PCR amplified DNA was determined for both strands by Sanger sequencing.

### MYXV Genome Sequencing

Purified viral DNA was first quantified using the Qubit dsDNA assay (Invitrogen) before being diluted to 0.2 ng/µl with molecular grade water. Sequence libraries were then prepared from the diluted viral DNA using the Nextera XT sample preparation kit (Illumina). Individual libraries were quantified using a KAPA library quantification kit (KAPA Biosystems) before manual normalization. The pooled libraries were then sequenced on an Illumina MiSeq using a 500 cycle v2 (250nt PE reads) at the Ramaciotti Centre for Genomics, Sydney with between 12 and 24 samples indexed in each run.

### Genome assembly

The sequence QC and genome assembly was performed using CLC Genomics v8 (Qiagen). First, the demultiplexed sequence reads were imported and quality trimmed to ensure removal of adapter sequences and low-quality bases (phred score < 20). Next, overlapping sequence reads were merged. The full set of trimmed sequence reads were then assembled *de novo* with a word (k-mer) size of 35. This typically yielded a large contig corresponding to the core genome (∼138K) and a small contig corresponding to the TIR (∼11K). The TIR contig was duplicated and one copy reverse complemented before manually assembling each onto the end of core genome (where sequence overlap was present) to generate a draft genome. Following this procedure, the full set of trimmed sequence reads was re-mapped back to check for polymorphisms, mis-alignments and the evenness of coverage, with the final majority consensus taken as the final genome assembly.

### Gene annotation

Genes were annotated as in the Lu reference (Brazil/Campinas-1949; NC001132) (20) and SLS (Brazil-1910; JX565574) (4) sequences. Gene numbers are italicized with the direction of transcription indicated as left (L) or right (R) (for example, *M010L*). Genes in the terminal inverted repeats are indicated by L/R. Proteins are indicated by gene number without the transcription direction (for example, M010). In the case of the *M156R* gene we revised the annotation to begin at a start codon downstream to the original annotation, such that the protein is 75 instead of 102 amino acids (46, 54).

### Evolutionary analysis

The 50 MYXV genome sequences determined here were combined with 25 previously sequenced viruses from the Australian outbreak. The European Lausanne sequence was used as an outgroup to root the tree. These sequences were initially aligned using MAFTT (75) and adjusted manually, resulting in a final sequence alignment data set of 78 MYXV sequences 163,711 bp in length and covering the years 1949-2016.

A phylogenetic tree of the aligned sequences was estimated using the maximum likelihood (ML) procedure available in the PhyML package (76). This analysis utilized the GTR+G_4_ model of nucleotide substitution and NNI+SPR branch-swapping. A bootstrap resampling procedure (1000 replications) was used to assess the statistical robustness of individual nodes on the MYXV phylogeny. To determine and visualize the extent of clock-like structure in these MYXV data we performed a regression of root-to-tip genetic distances against year of sampling on the ML tree using TempEst (77).

To test for the presence of recombination in the Australian MYXV data we first performed a screen for recombination using the RDP, Genecov and Bootscan methods (with default settings) available within the RDP4 package (78). In addition, we estimated ML phylogenetic trees for non-overlapping 25K sections of all the Australian MXYV sequences sampled here using the same procedure as described above. A visual inspection for phylogenetic incongruence supported by high numbers of bootstrap replications (>70%) was then used to identify putative recombination events.

### GenBank accession numbers

All MYXV sequences generated here have been submitted to GenBank and assigned accession numbers XXX-YYY.

## Acknowledgements

Rabbit samples were kindly provided by Dr Tarnya Cox and Dr Pam Whitely. Mrs Karen Glover provided technical support. ECH is funded by an ARC Australian Laureate Fellowship (FL170100022). This work was funded in part by the National Institute of Allergy and Infectious Diseases (grant R01AI093804)

